# Marked compositional changes in harvestmen assemblages in Amazonian forest islands induced by a mega dam

**DOI:** 10.1101/542969

**Authors:** Ana Lúcia Tourinho, Maíra Benchimol, Willians Porto, Carlos A. Peres, Danielle Storck-Tonon

**Affiliations:** Programa de Pós-Graduação em Ciências Ambientais, Universidade Federal do Mato Grosso, Instituto de Ciências Naturais, Humanas e Sociais, Núcleo de Estudos da Biodiversidade da Amazônia Mato-Grossense (NEBAM), Av. Alexandre Ferronato, 1200, Setor Industrial, CEP: 78577-267, Sinop, MT, Brazil; Laboratório de Ecologia Aplicada à Conservação, Universidade Estadual de Santa Cruz, Rodovia Jorge Amado km 16, CEP: 45662-900 Ilhéus, BA, Brazil; División de Aracnología, Museo Argentino de Ciencias Naturales–CONICET, Av. Ángel Gallardo 470, C1405DJR Buenos Aires, Argentina; School Environmental Sciences, University of East Anglia, Norwich NR47TJ, UK; Departamento de Sistemática e Ecologia, Universidade Federal da Paraíba, João Pessoa, CEP 58051-900, Brazil; Programa de Pós-Graduação em Ambiente e Sistemas de Produção Agrícola, Universidade do Estado de Mato Grosso, Rod. MT 358, km 7 - Jardim Aeroporto. CEP: 78300-000, Tangará da Serra, MT, Brazil

**Keywords:** Conservation biology, landscape ecology, habitat fragmentation, hydroelectric dams, environmental quality, indicator species

## Abstract

1. Mega hydroelectric dams have become one of the main drivers of habitat loss in tropical forests, converting large tracts of pristine forests into isolated forest islands. Understanding how biodiversity cope with landscape modification in these archipelagic landscapes is of paramount importance to assess the environmental consequences of dam infrastructure and propose mitigation actions for biodiversity conservation. In this context, harvestmen (Opiliones, Arachnida) comprise an excellent indicator taxon of habitat quality, given their high sensitivity to desiccation and microclimatic change.
2. We investigate the effects of landscape change induced by a mega hydropower dam on forest harvestmen species richness, abundance and composition within the Balbina Hydroelectric Dam, Central Brazilian Amazon. We sampled 20 islands and five mainland continuous forests, relating our biological response variables to local, patch and landscape scale metrics.
3. Although unexpectedly species richness was unaffected by any local, patch and landscape variables, species composition and abundance were differentially affected by a set of predictor variables at different scales. Forest cover and fallen woody stems were significant predictors of patterns of species composition, whereas vegetation density, forest cover, island area, abundance of palm trees, and fallen woody stems best explained harvestmen abundance.
4. Our results indicate that both islands embedded within greater and lower amount of forest cover are important to ensure high diversity of harvestmen. We recommend retaining large forest habitat patches surrounded by a large amount of forest cover to minimize forest disturbance effects and enhance long-term persistence of harvestmen sensitive species in large hydroelectric dams.

## Introduction

Hydroelectric dams have been posed as a major new villain for Amazonian rainforest and freshwater ecosystems, severely inducing the widespread habitat loss and wholesale local species extinctions (Benchimol & Peres, 2015a; Bobrowiec & Tavares, 2017, Lees *et al*., 2016). The Amazon basin accounts for 18% of global scale river discharge (Dai & Trenberth, 2002), so it is not surprising that this region has rapidly become a major target for hydropower development; in addition to 191 existing dams, 246 additional dams may be built across the whole Amazon Basin in the next few decades (Lees *et al*. 2016). Indeed, more than 1,300,00 ha of forest has already been flooded by operational dams in the Brazilian Amazon alone (ECOA, 2018).

Although mega-dam infrastructure projects have often been referred to as ‘green energy’ sources (renewable energy resources that provide the highest environmental benefits) (Gibson *et al*., 2017), several studies have shown the pervasive social and environmental impacts of large dams. From local communities (Esselman & Opperman, 2010, Fearnside, 2016) to fluvial hydrology (Nilsson *et al*., 2005), large dams affect indigenous communities by flooding their territories, and increasing greenhouse gas emissions (Almeida *et al*., 2010). They also drive local extinctions of both aquatic (Alho, 2011; Liermann *et al*., 2012; Palmeirim *et al*., 2014) and terrestrial fauna and flora (Terborgh *et al*., 1997; Benchimol & Venticinque, 2014, Benchimol & Peres, 2015a,b,c). However, few studies have assessed the effects of large dams on terrestrial invertebrates, despite their immense importance for forest nutrient cycling and ecosystem services (Storck-Tonon & Peres, 2017, Carvalho *et al*., 2012). For instance, a study has shown that most orchid bee species are driven to local extinction after 26 years of insularization within a mega dam in central Brazilian Amazonia, with potential effects on pollination (Storck-Tonon & Peres, 2017).

Several groups of arthropods can be described as effective bioindicators, providing valuable information on environmental quality at both small and large scales on the basis of rapid surveys, enabling their efficient inclusion in environmental licensing studies prior to development projects (Maleque *et al*., 2006, Melo *et al*., 2015). Additionally, arthropods play an important role in many key processes influencing ecosystem cycling of carbon and other nutrients, and modulating the amount and quality of resources entering the detrital food web (Yang & Gratton, 2014). These processes may occur via direct or indirect pathways mediated through species interactions via plants and soil microbes, integrating community-level interactions with ecosystem processes. Arachnids are mostly generalist predators, and are often more dependent on the physical structure of microhabitats than on prey availability (Villanueva-Bonilla *et al*., 2017). Most harvestmen species (Opiliones, Arachnida) inhabit humid habitats; indeed, they are physiologically dependent of humid environments, due to their high susceptibility to desiccation, nocturnal activity and low vagility (Pinto-da-Rocha *et al*., 2007).

Despite the significant ecological importance of harvestmen (Fig. 2), serving as excellent predictors of environmental quality given their rapid responses to habitat change and fragmentation (Bragagnolo *et al*., 2007, Černecká *et al*., 2017), this charismatic taxonomic group remains poorly investigated in tropical fragmented forest landscapes. Considering that fragmentation is a landscape-scale process (Fahrig, 2003), studies assessing the responses of harvestmen communities to forest fragmentation should explicitly consider the influence of landscape variables. Under the island biogeography paradigm (MacArthur & Wilson, 1967), fragment size and isolation are frequently hailed as primary determinants of species richness in remaining habitat patches. Yet several studies have clearly emphasized the importance of also considering the spatial arrangement of forest fragments and habitat quality. For instance, the percentage of surrounding habitat within the landscape has been widely considered as a good predictor of species diversity for several vertebrate and floristic groups (Andrén, 1994; Morante-Filho *et al*., 2015; Benchimol *et al*., 2017), and the dominant role of habitat amount has been suggested when species richness is measured at sample sites (Fahrig, 2013; but see Bueno & Peres 2019). Thus, combining different patch and landscape metrics are likely to provide a better understanding on the effects of habitat fragmentation on the diversity patterns of harvestmen assemblages.

Additionally, patterns of diversity in harvestmen assemblages are highly correlated with vegetation structure at meso and local scales, and are especially responsive to the presence of large old trees and palms in Amazonian forests (Colmenares *et al*., 2016; Tourinho *et al*., 2014, 2018, Porto *et al*., 2016). This occurs because palms increase habitat complexity at ground level, as a consequence of the fallen litter trapped within their leaves (Vasconcelos, 1990, Franken *et al*., 2013). Therefore, both palm and non-palm trees are directly responsible for the quality, depth and decomposition of the soil leaf-litter layer, which are a key microhabitat components affecting assemblages of small-bodied harvestmen species and other invertebrates (Tarli *et al*., 2014, Colmenares *et al*., 2016, Porto *et al*. 2016).

Although few studies have shown the negative impacts of fragmented forest habitats on harvestmen assemblages in temperate forests (Černecká *et al*., 2017, Mihál & Černecká, 2017), only one study to date -- conducted in forest patches embedded within heterogeneous vegetation matrices in the Brazilian Atlantic forest -- has assessed the effects of forest fragmentation on harvestmen within a tropical forest landscape (Bragagnolo, *et al*. 2007). In this study, harvestmen assemblages were more clearly structured by forest quality and size than other arthropod taxa, suggesting that they are good community-wide indicators of intermediate-level disturbances and can be used as a robust model to monitor tropical forest disturbance. However, we still lack information on how this key taxonomic group is affected by habitat insularization, with archipelagic ecosystems formed in the aftermath of rising floodwaters induced by large dams providing a unique natural experiment in this context.

Here, we assessed the degree to which forest fragmentation induced by a mega hydroelectric dam affects harvestmen assemblages in the Brazilian Amazon. We conducted sampling surveys at five mainland continuous forest sites and 20 widely distributed forest islands differing in size, shape and degree of isolation within the ∼443,772-hectares Balbina Hydroelectric Dam. Specifically, we investigate (1) the overall patterns of harvestmen structure in insular forest environments; and (2) the potential effect of landscape features and habitat structure on harvestmen assemblages. For this, we firstly investigated the area and isolation effects following the equilibrium theory of island biogeography, and then examined the degree to which patch and landscape metrics and local habitat variables can predict harvestmen species richness, abundance and composition across forest islands.

## Materials and Methods

### Study site

We conducted this research at the Balbina Hydroelectric Reservoir landscape - BHR (1°48’S; 59°29’W), located in the state of Amazonas, central Brazilian Amazonia. The Balbina Dam is a large man-made reservoir that was formed in 1987 by the damming of the Uatumã river, aiming to supply power to the growing capital city of Manaus. An area of 312,900 ha was inundated, resulting in the creation of 3,546 islands ranging in size from <1 to 4,878 ha. Most islands are covered by primary sub-montane dense forests, and are surrounded by open-water and dead trees that rise above the water level; the mean water column depth range from 7.4 m (in most of the reservoir) to 30 m (in the former river channel; Eletronorte, 1997). Mean annual temperature is 28°C and the mean annual precipitation is 2376 mm. A large tract of pristine continuous forest sites surrounds the Balbina archipelagic landscape, and share similar floristic composition, fauna, climate and topography (Benchimol & Peres, 2015a,c). In 1990, part of the archipelago and an extensive tract of neighbouring pristine forest became protected by the creation of the ∼940,358-ha Uatumã Biological Reserve, the largest Biological Reserve in Brazil. Although islands are unaffected by logging and hunting pressure, several islands experienced ephemeral understorey fires during the El Niño drought of late-1997 (Benchimol & Peres, 2015a).

### Study Design

We selected 20 islands using two cloudless georeferenced Landsat ETM+ scenes. Additionally, we surveyed five continuous (‘pseudo-control’) forest sites located in the southernmost part of the Uatumã Reserve, within a permanent study grid of the PPBio (Brazilian Biodiversity Research Program, Fig. 1). Islands were selected on the basis of their size, shape, isolation and spatial distribution, keeping a minimum distance of 1 km from one another (Fig. 1). Likewise, each continuous forest site in the mainland was also spaced by 1 km from each other. Island area ranged from 11.8 to 218.5 ha (mean ± SD = 91.5 ± 69.7 ha), and ranged widely in isolation distances [to the nearest mainland continuous forest] from 10 m to 9,000 m (mean ± SD = 3,220 ± 2,648 m).

**Figure 1.**
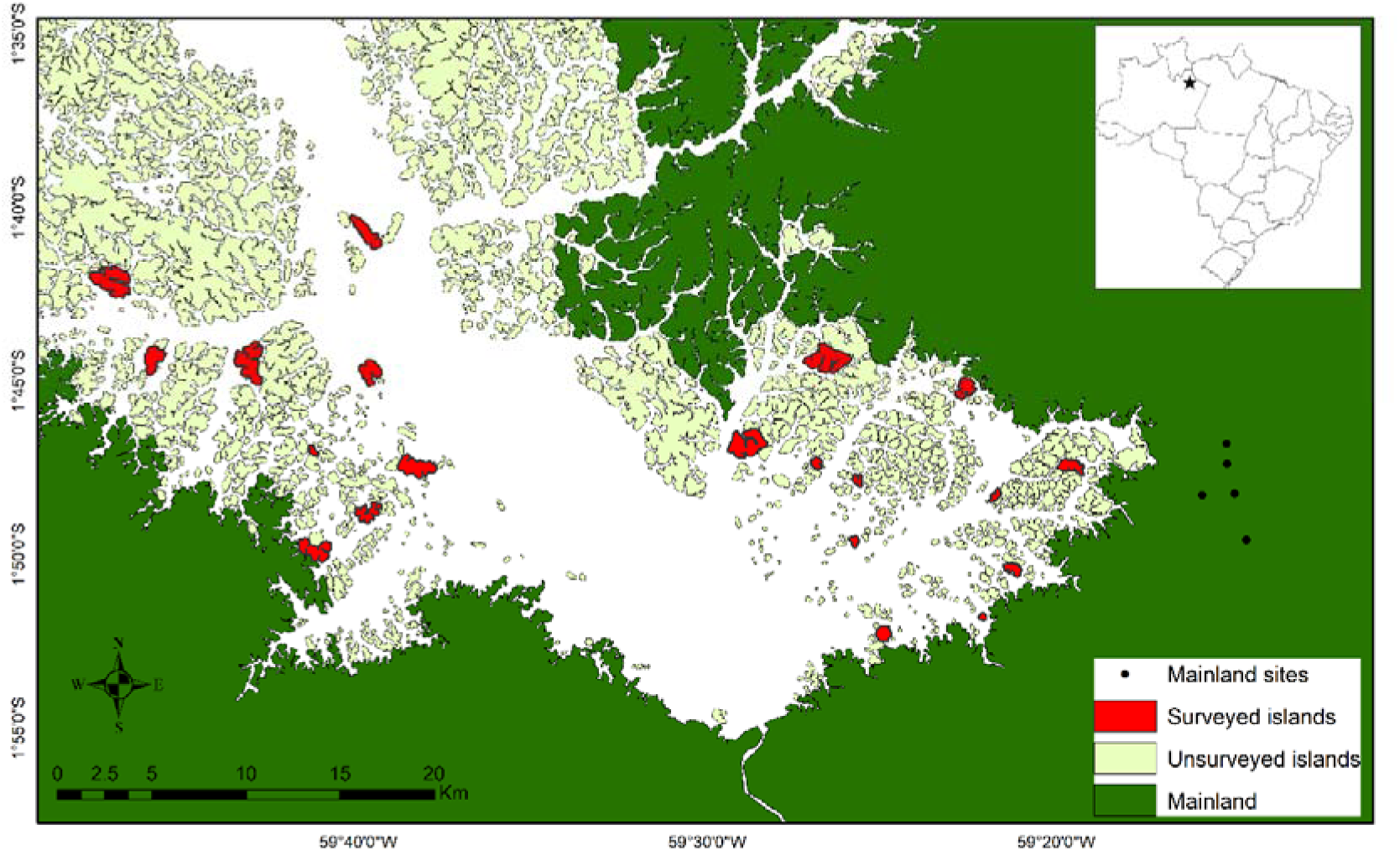
Spatial distribution of the 20 sampled islands (red) and 5 continuous forest sites (black circles) within the Balbina Hydroelectric Reservoir of central Brazilian Amazonia.

**Figure 2.**
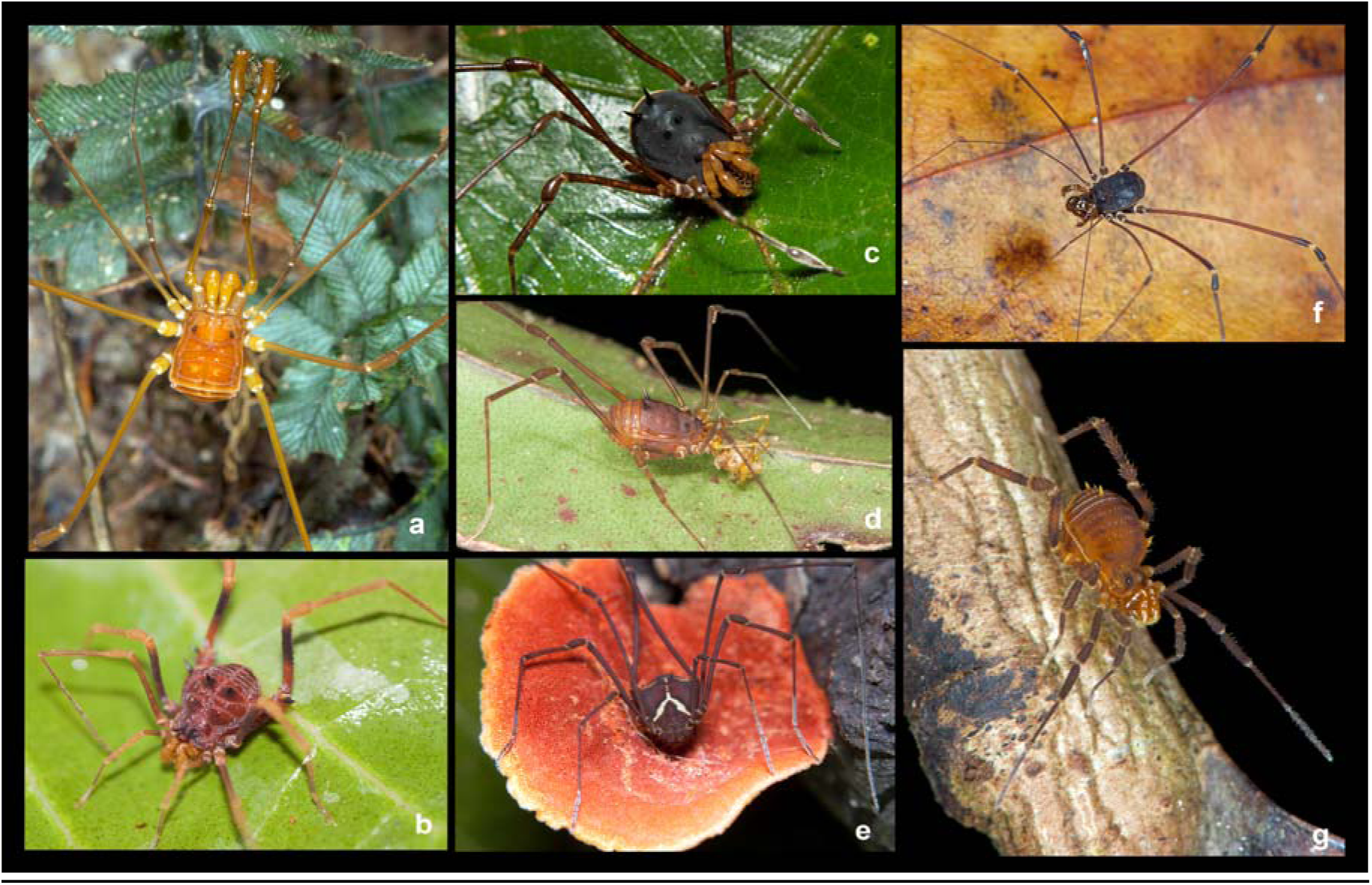
Some of the harvestmen species occurring in the Balbina Hydroelectric Reservoir - BHR and mainland: a) Male of *Protimesius longipalpis* (Stygnidae); b) Female of *Amazochroma carvalhoi* (Gonyleptidae), c) Male of *Saramacia lucaseae* (Manaosbiidae); d) Female of *Stygnus pectinipes* (Stygnidae); e) Male of *Eucynortella aff. duapunctata* (Cosmetidae); f) Male of *Avima matintaperera* (Agoristenidaae); g) Male of *Fissiphallius martensi* (Fissiphalliidae).

### Harvestmen sampling

Between 2006 and 2008, we systematically sampled harvestmen assemblages (Fig. 2) using four 300-m^2^ transects (30 m × 10 m) at each forest site, separated by 30 m, totalling 1,200 m^2^ of transect area per site (see Tourinho *et al*., 2014, 2018, Porto *et al*., 2016). To avoid edge effects on islands, transects were established at least 250 m from the nearest forest edge. Each plot was surveyed using both an active nocturnal search method (Sørensen *et al*., 2002; Pinto-da-Rocha & Bonaldo, 2006; Resende *et al*., 2012) and diurnal beating tray surveys. Nocturnal searchers were performed by one collector for 1 hour, during which all harvestmen (and other arachnids) encountered on the forest floor to a height of 2 m, on tree trunks and other vegetation were captured. Beating tray surveys were conducted during daylight hours, by one collector for 1 h, during which shrubs up to 3 m in height were struck 20 times with a wooden stick. We used a 1-m^2^ white fabric wooden frame positioned underneath the shrubs to capture any fallen harvestmen.

All individuals collected were subsequently identified at the lowest taxonomic level by experts on harvestmen taxonomy (Ana Lúcia Tourinho, Pío Colmenares & Willians Porto). For this, we examined the external morphology of individual specimens under a stereomicroscope, which were compared with original descriptions provided in the literature (Pinto-da-Rocha, 1994, 1996, 1997, 2004; Kury, 2003), type specimens, and photographs of type specimens. We excluded from the dataset nymphs and females, which have ambiguous morphology, because they cannot be assigned to any genus or species level. In the case of species groups with a very conservative external morphology and/or poorly understood taxonomy (e.g. Cosmetidae, Sclerosomatidae, Zalmoxidae) we also examined their male genitalia to improve species delimitation, following Acosta *et al*. (2007). All specimens collected were preserved in 70% ethanol and deposited in the Invertebrate Collection of the Instituto Nacional de Pesquisas da Amazônia, Manaus, Amazonas, Brazil, under curator, Dr Célio Magalhães.

### Landscape and habitat variables

We used RapidEye© high-resolution (5-m pixel) imagery for the entire Balbina archipelago to quantify landscape structure variables at the patch and landscape-scales, associated with each of surveyed forest site. At the patch scale, we obtained the total island area in hectares (log_10_ x; ‘area’); the isolation distance from each island to the nearest mainland (‘*D*_mainland_’); and the island shape (‘shape’) by dividing the total island area by the total island perimeter using ArcMap (ESRI, 2016). At the landscape scale, we estimated the total amount of forest cover (‘cover’) within a specified radius, and modified the proximity index of McGarigal *et al*., (2012) by considering the total size of any land mass within the buffer, rather than excluding land areas outside the buffer within patches encompassed by the buffer (‘prox’). We considered multiple buffers (250m, 500m and 1000m) outside the perimeter of each focal island and mainland forest sites for landscape-scale analyses.

We selected variables at local scale that were previously documented and/or tested to influence harvestmen assemblages. In this regard it was demonstrated that harvestmen assemblages use trees as refuges when disturbed, but only in sites with higher harvestmen diversity in the ground/litter microhabitat (Proud et al., 2012). Recent research also offered evidence for a relationship between number of palm trees and harvestmen assemblage composition in upland Amazonian forest (Tourinho et al., 2014; Colmenares et al., 2016; Porto et al., 2016). In previous studies in Amazon upland forests, most abundant harvestmen species were found mostly in trees (Colmenares Tourinho et al., 2014; Porto et al., 2016), being especially common in the accumulated litter at the base of the fronds in stemless and arborescent palms. Fallen logs, roots, termite nests, and suspended litter at the base of trees and palm leaves are the habitats favored by harvestmen, providing ideal microclimatic conditions of humidity and temperature for their development (Mestre & Pinto-da-Rocha, 2004; Curtis & Machado, 2007; Curtis, 2007; Proud et al., 2012). We also obtained at local scale the following variables the litter depth (‘litdepth’), the number of trees (‘trees’), the total number of palm trees (‘palms’) and the number of fallen woody stems (‘fallen’). ‘Fallen’ considered down wood and does not include arborescent palm left bracts all of which were used here as proxies of vegetation habitat structure available to harvestmen (Colmenares *et al*. 2016). For this, we mapped all non-palm and palm trees with a diameter at breast height (DBH) > 30 cm within each 300-m^2^ transect (Table 1). We also mapped all fallen stems within each plot. At every 5 m along the long axis of the transects, a measurement of litter depth was taken. These measurements consisted of forcing a wooden stick 0.5 cm in diameter into the litter until it reached the soil and noting the height (in cm) above ground to the top layer of litter.

**Table 1.**
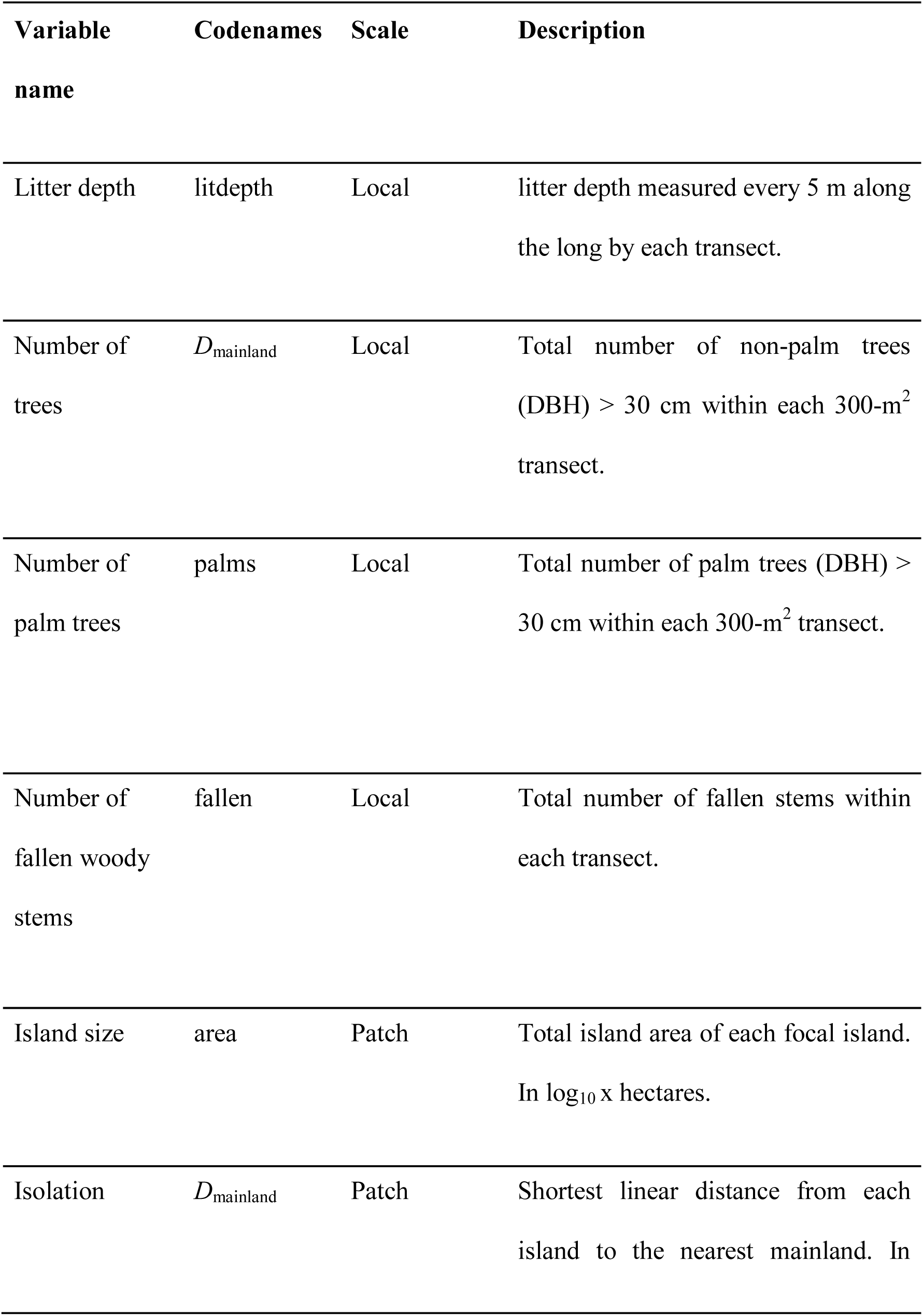
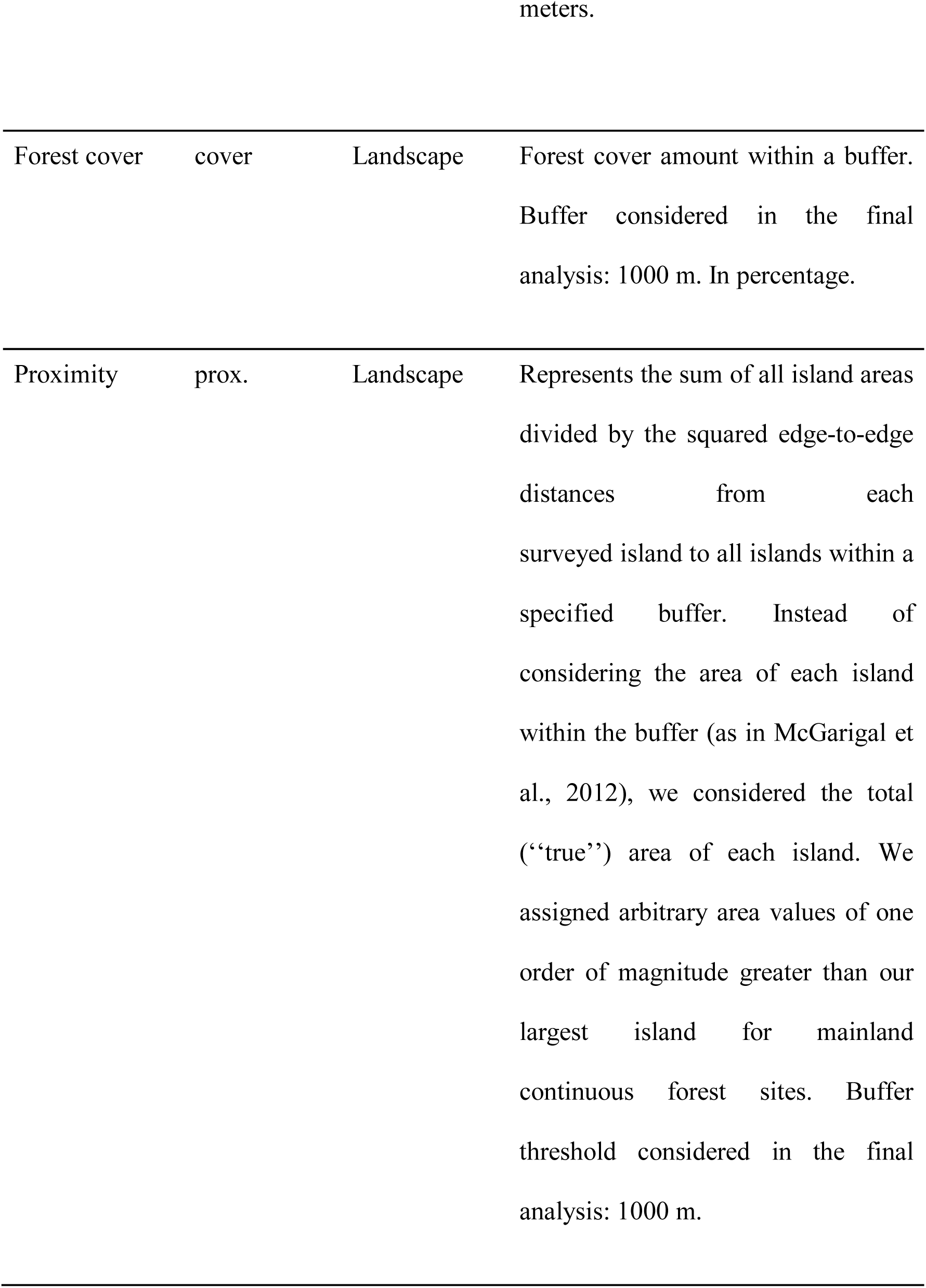
Description of the explanatory variables used in the analyses.

### Data analysis

We pooled data from the four sampled transects on each island and each mainland site, each of which were used as a sampling unit. We firstly compared the coefficient of determination (R^2^) from regression analysis of each community attribute investigated (i.e., species richness, overall abundance and species composition) for each forest cover radius (i.e., 250 m, 500 m, 1000 m) to determine which radial buffer best explained diversity patterns. The highest R^2^ values were obtained at 1000 m radius (see Table S1), so we used this buffer size for all subsequently analyses. We thus performed a Pearson correlation test using all landscape and habitat variables, excluding variables that were highly correlated (r ≥ 0.50) from the analysis, resulting in five habitat and landscape variables maintained in the global models: area, litter depth, number of palm trees, number of fallen woody stems, and forest cover. Among the correlated variables we have chosen the variables that were previously documented and/or tested to have a strong influence in harvestmen assemblages.

To assess differences in harvestmen assemblage structure among the habitat types (islands and mainland) we performed permutational multivariate analysis of variance (PERMANOVA) using the landscape and habitat variables. We used a multiple regression on distance matrices analysis (MRM) to analyse changes in species composition and test for spatial correlation between all surveyed sites, this analysis is similar to partial Mantel tests and was used to examine the levels of correlation and the spatial independence between the dependent distance matrix and the independent explanatory distance matrices. For that we used the abundance matrix of each species obtained within the transects, then the longitude and latitude matrix to create a geographic distance matrix and all the landscape and habitat variables cited above to create the explanatory variables distance matrices that were used to perform the MRM.

Following the theory of island biogeography (MacArthur & Wilson 1967), we first examined the potential effects of island size (log(x+1)) and isolation (*D*_mainland_) on harvestmen species richness, abundance and composition using linear regression. For the latter, we reduced the data dimensionality using NMDS ordinations based on two axes, considering both the Jaccard (presence/absence) and Bray-Curtis (abundance) similarity values, and used the first axis as a predictor of species composition.

We further performed Generalized Linear Models (GLMs) to examine the relationship between harvestmen species richness, abundance and composition (NMDS1) of islands and the explanatory variables. Multicollinearity among these variables was tested using Variation Inflation Factors (VIF) (Domman *et al*. 2013) using the ‘HH’ package (Heiberger, 2016), deleting the least moderately redundant or collinear variables (VIF > 6). We used Poisson, Gaussian and quasi-Poisson error structures for GLMs of species richness, composition, and abundance, respectively. For all response variables, the global model included all non-collinear environmental variables. We thus ran all possible combinations of subsets in addition to the null model using the ‘MuMIn’ package (Barton, 2018) and retained all models that differed by ΔAIC ≤ 2.00 (Burnham & Anderson 2002). All analyses were performed within the R 3.3.1 environment (R Core Team 2016) and RStudio Team (2016).

## Results

A total of 783 harvestmen representing 31 species, 22 genera and 9 families were collected across all sampling sites throughout the BHR (Table 3). These ranged along the size spectrum from the small-bodied Fissiphalliidae, Zalmoxidae and Stygnomatidae (2.00 ± 2.50 mm) to the large-bodied Stygnidae species of *Protimesius* (4.95 ± 10.85 mm) (Fig. 2). The number of species co-occurring within any given island or continuous forest site ranged from 1 to 20 (mean ± SD = 6.25 ± 4.14) species. The families containing the largest numbers of species recorded were Stygnidae (9), Sclerosomatidae (6), Manaosbiidae (4), and Cosmetidae (4) (Table 3), whereas the most abundant families were Stygnidae (209) and Manaosbiidae (202). Nine species were collected exclusively on islands, 14 were exclusively collected at mainland sites, and nine in both islands and mainland sites (Fig. 5, Table 3). The families with largest number of species on islands were Stygnidae (7) and Zalmoxidae (3), whereas the Sclerosomatidae (6) and Stygnidae (5) were the most speciouse families in the mainland. All Sclerosomatidae (194 individuals, 6 species) were exclusively restricted to mainland sites (Fig. 5). The number of species per island was much lower than recorded at mainland forest sites. Indeed, 55% of the surveyed islands exhibited only one to five species, 45% harboured more than five species, and three islands contained only 7, 8 and 10 species; in contrast, continuous forest sites in the mainland retained between 11 and 20 species.

Considering only islands, there was no significant species-area relationship (SAR) on the basis of a semi-log model (R^2^_adj_ = 0.000, P = 0.471). Island distance to the nearest mainland area also failed to explain harvestmen species richness (R^2^_adj_ = 0.000, P = 0.835). Similar patterns were also observed for species composition, with neither island size (R^2^_adj_ = 0.001, P = 0.326) nor island isolation (R^2^_adj_= 0.000, P = 0.422) significantly affecting the first NMDS axis. Island size and isolation also failed to affect patterns of overall harvestmen abundance within sites (Area - R^2^_adj_ = 0.000, P = 0.423; Isolation - R^2^_adj_ = 0.000, P = 0.363). The species composition of harvestmen (Fig. 3) differed between islands and mainland continuous forest sites (PERMANOVA: F= 4.76, P =0.01). None of compositional metrics were affected by distance (MRM r = 0.24, p = 0.180).

**Figure 3.**
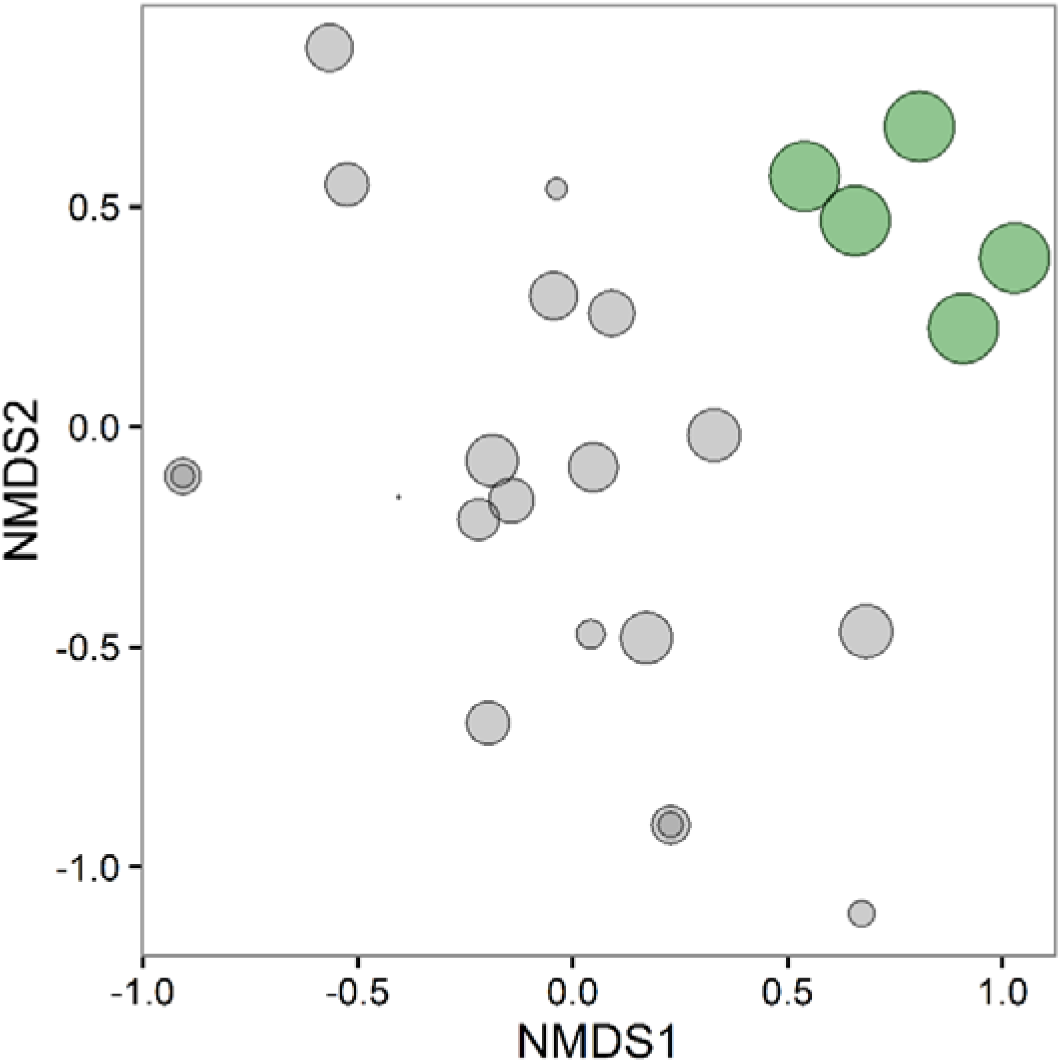
Nonmetric multidimensional scaling (NMDS) ordination using a Jaccard similarity matrix of harvestman composition in islands (gray circles) and mainland forest (green circles) sampled across the Balbina archipelago. Circles are sized in proportion to the log area of sampled sites.

When all explanatory variables were also incorporated into the models, GLMs showed that landscape-scale forest cover alone was included in the best model explaining patterns of local species composition (Table 2; Fig. 4). Considering species richness, however, neither of the local habitat nor landscape variables were significant predictors in the ‘best’ model. In contrast, number of trees, forest cover, island size, abundance of palm trees, and the amount of fallen woody stems were all significant predictors of harvestmen abundance (Table 2).

**Figure 4.**
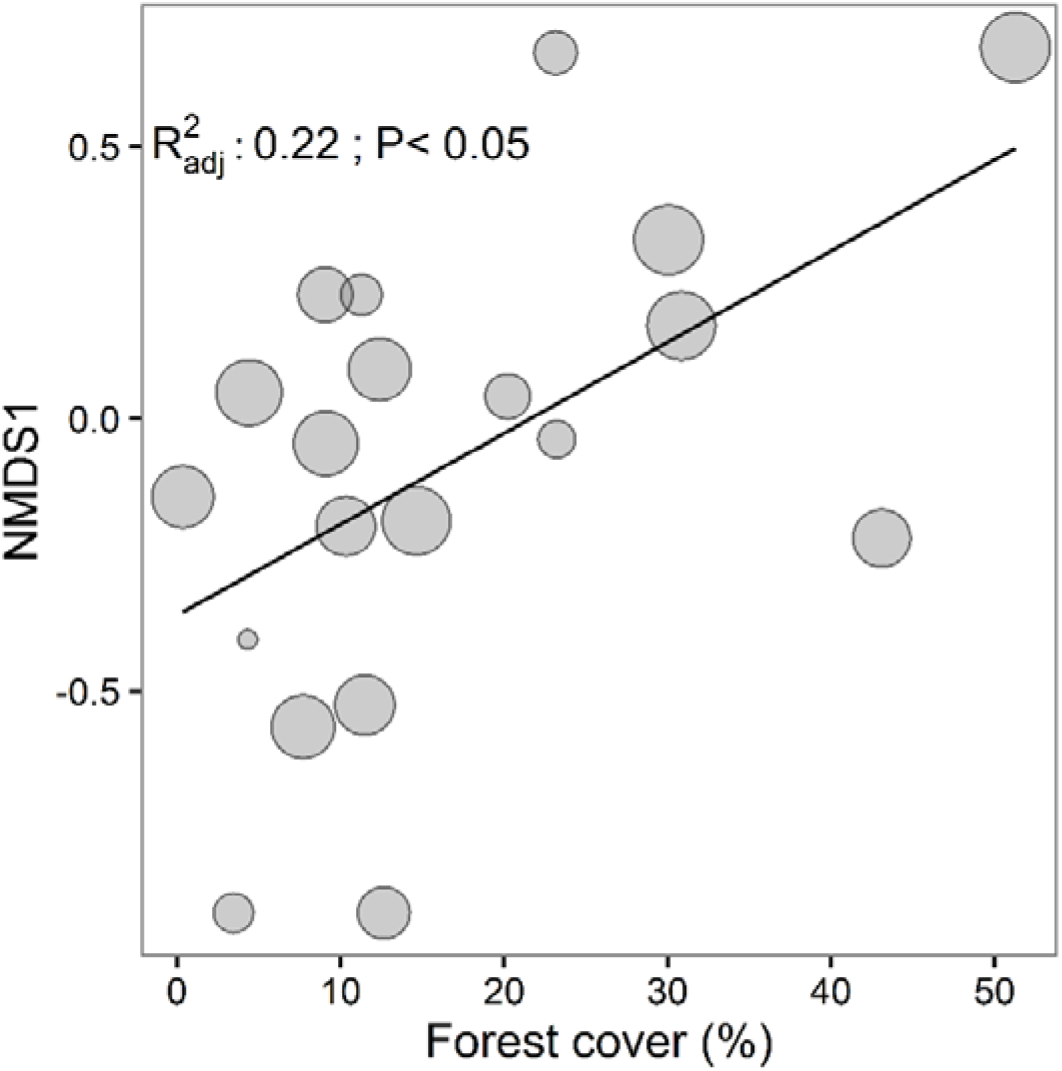
Relationships between the first non-metric multidimensional scaling (NMDS) axis representing the harvestmen species composition and forest cover within a 1000 m buffer. Circles proportional to the log area.

**Figure 5.**
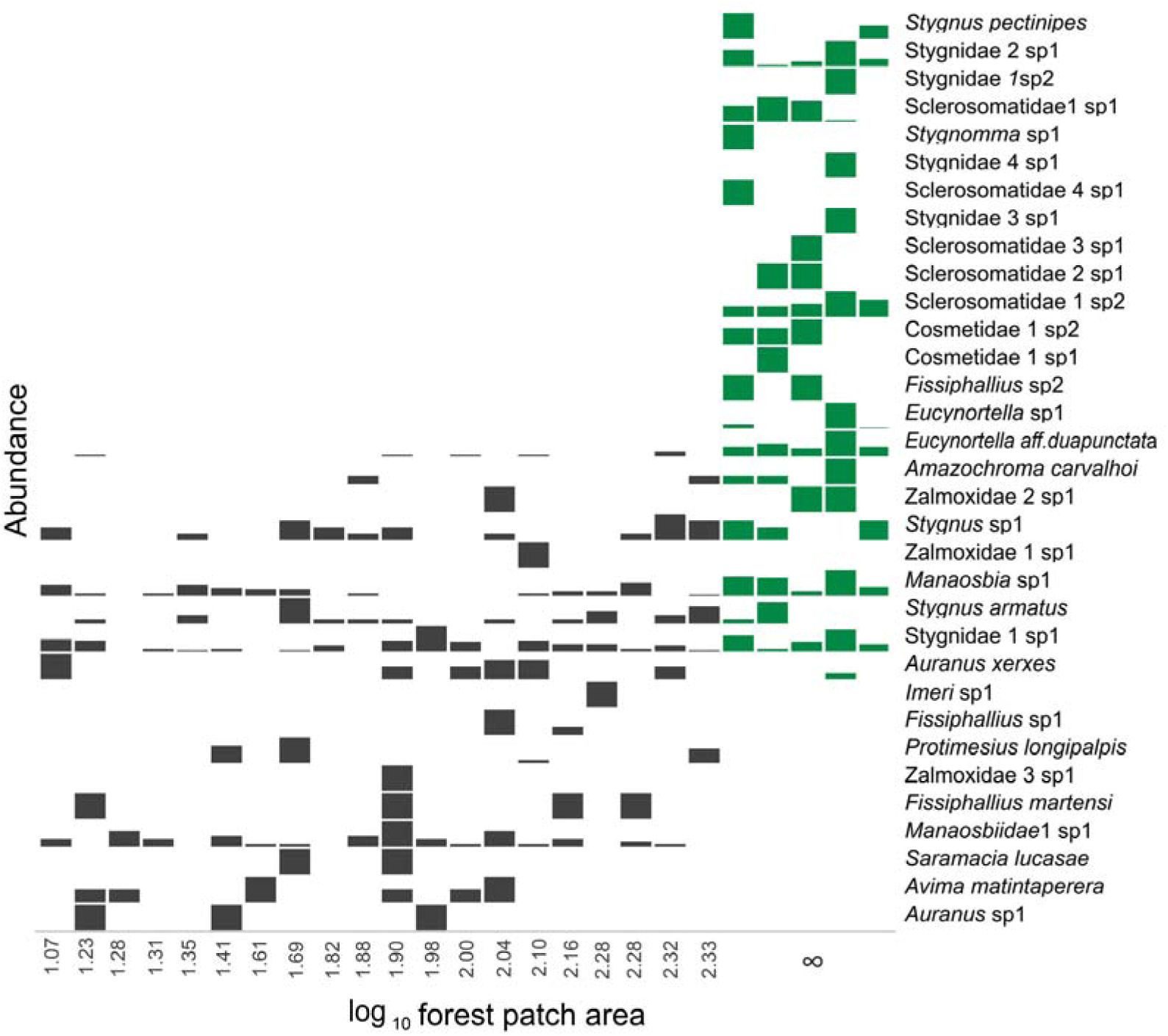
Site records of 31 harvestman species sampled across 25 trapping sites distributed throughout the Balbina Hydroelectric Reservoir. Species abundances within island and mainland forest sites are represented by gray and green rectangles, respectively.

**Table 2.** Results of the Generalized Linear Models for the effects of island area (log10), litter depth, fallen woody stems, palm trees, forest cover (buffer 1000m) and trees. Only the best models are shown (ΔAICc <2).

| Response variables | Model | Intercept | Area | Forest cover | Fallen<br>woody stems | Trees | Palm<br>trees | Litter<br>depht | AICc | ΔAICc | Akaike<br>weights |
| --- | --- | --- | --- | --- | --- | --- | --- | --- | --- | --- | --- |
| Species Richness | 1 | 1.589 |  |  |  |  |  |  | 88.7 | 0.00 | 0.116 |
|  | 17 | 1.155 |  |  |  |  |  | 0.171 | 89.4 | 0.77 | 0.070 |
| Species composition | 3 | -0.306 |  | 0.004 |  |  |  |  | 23.8 | 0.00 | 0.244 |
|  | 35 | -0.524 |  | 0.004 | 0.061 |  |  |  | 25.4 | 1.60 | 0.109 |
| Species abundance | 48 | 1.910 | 1.109 | -0.014 | -0.090 | -0.288 | 0.018 |  | 220.7 | 0.00 | 0.253 |
|  | 32 | 1.918 | 0.594 | -0.01 |  | -0.281 | 0.017 | 0.209 | 220.8 | 0.10 | 0.241 |

**Table 3.** Number of individuals of each harvestmen species recorded at the Balbina Hydroelectric Reservoir.

| Species | Mainland | Island | Total |
| --- | --- | --- | --- |
| AGORISTENIDAE |  |  |  |
| *Avima matintaperera* (Pinto-da-Rocha, 1996) |  | 8 | 8 |
| COSMETIDAE |  |  |  |
| *Cosmetidae1* sp.1 | 42 |  | 42 |
| *Cosmetidae1* sp.2 | 13 |  | 13 |
| *Eucynortella* aff. *duapunctata* Goodnight & Goodnight, 1943 | 41 | 7 | 48 |
| *Eucynortella* sp.1 | 44 |  | 44 |
| FISSIPHALLIIDAE |  |  |  |
| *Fissiphallius martensi* Pinto-da-Rocha, 2004 |  | 4 | 4 |
| *Fissiphallius* sp.1 |  | 4 | 4 |
| *Fissiphallius* sp.2 | 2 |  | 2 |
| GONYLEPTIDAE |  |  |  |
| *Amazochroma carvalhoi* (Mello-Leitão, 1941) | 5 | 2 | 7 |
| MANAOSBIIDAE |  |  |  |
| *Manaosbia* sp.1 | 17 | 120 | 137 |
| *Manaosbiidae1* sp.1 | 4 | 58 | 62 |
| *Manaosbiidae2* sp.1 | 2 |  | 2 |
| *Saramacia lucasae* Jim & Soares, 1991 | 1 |  | 1 |
| SCLEROSOMATIDAE |  |  |  |
| *Sclerosomatidae1* sp.1 | 48 |  | 48 |
| *Sclerosomatidae1* sp.2 | 58 |  | 58 |
| *Sclerosomatidae2* sp.1 | 79 |  | 79 |
| *Sclerosomatidae2* sp.2 | 2 |  | 2 |
| *Sclerosomatidae3* sp.1 | 4 |  | 4 |
| *Sclerosomatidae4* sp.1 | 3 |  | 3 |
| STYGNIDAE |  | 3 | 3 |
| *Auranus* sp.1 |  |  |  |
| *Auranus xerxes* Colmenares, Porto & Tourinho, 2016 | 1 | 16 | 17 |
| *Imeri* sp.1 |  | 2 | 2 |
| *Protimesius longipalpis* Roewer, 1943 | 23 |  | 23 |
| *Stygnidae1* sp.1 | 12 | 88 | 100 |
| *Stygnus armatus* Perty, 1833 | 14 | 18 | 32 |
| *Stygnus pectinipes* (Roewer, 1943) |  | 3 | 3 |
| *Stygnus* sp.1 | 9 | 20 | 29 |
| STYGNOMMATIDAE |  |  |  |
| *Stygnomma* sp.1 |  | 1 | 1 |
| ZALMOXIDAE |  |  |  |
| *Zalmoxidae1* sp.1 |  | 1 | 1 |
| *Zalmoxidae2* sp.1 | 2 | 1 | 3 |
| *Zalmoxidae3* sp.1 |  | 1 | 1 |
| Total | 426 | 357 | 783 |

## Discussion

As far as we are aware, this is the first study to examine harvestmen responses to habitat insularization induced by a large dam within tropical forests. Islands harboured fewer species than mainland continuous forest sites, and exhibited a simplified species composition. Contrary to our expectations, island area and isolation did not predict patterns of species richness, as predicted by the equilibrium theory of island biogeography (MacArthur & Wilson, 1967). However, a set of local, patch and landscape variables was included in the best models predicting species composition and abundance. Due to the detrimental impacts of hydroelectric dams on forest biota, we provide additional evidence that these infrastructure projects are far from the ‘green energy’ sources they are often considered to be (Gibson *et al*., 2017), and propose conservation actions to enhance the long term persistence of harvestmen assemblages and other arthropods within reservoir areas.

### Main drivers of harvestmen assemblage structure

Contrary to our expectations, none of the local, patch and landscape variables were included in the best models explaining harvestmen species richness on islands. In the Balbina archipelagic landscape, island size was a strong positive predictor of species richness in other taxonomic groups including birds (Aurélio-Silva *et al*., 2016), small ─ mammals (Palmeirim *et al*. 2018) and large mammals (Benchimol & Peres, 2015a). Although the species-area relationship is one of the few ironclad rules in ecology (Mathews *et al*. 2016), we found that area failed in explaining richness patterns for harvestmen assemblages, the absence of this relationship has been documented before for this group in Amazonian upland forest sites where microhabitat structure and availability per unit area also explained patterns of harvestmen species composition but not species richness in harvestmen assemblages (Colmenares et al., 2016). The same pattern was also found for other arthropods, such as macro-moth communities in temperate woodland countryside landscapes, although in that case habitat amount, rather than patch size and isolation were the main drivers of species richness (Merckx *et al*., 2019), both however have not explained it for harvestmen either. Harvestmen species richness may not be responding to any of the most common predictor variable collectively but individually some severe richness reduction occurred in the fragmented Island mosaic in Balbina lake, on the other hand there are some species apparently benefiting of the fragmentation effects promoted by the dam.

Although the species area relationship is one of the most known and well documented patterns in ecology our results shown that considering SAR as more important or only considering SAR may possibly mislead or obfuscate the main conclusions of the impacts of forest fragmented areas. There were marked shifts in species composition, and we detected a significant species turnover between islands and continuous forest sites, indicating pervasive changes of harvestmen species assemblages on islands after 20 years of isolation. Because the continuous forest sites are clustered far from the islands (Fig. 4), it is possible that some differences in species composition were simply caused by the geographical separation of these areas. However, it is unlikely that such a strong difference in species composition as shown (Fig. 5) could be entirely caused by distance-decay. Both forest cover and fallen woody stems were the main joint drivers structuring the composition of harvestmen species on islands. Given that the presence of large canopy trees and palm tree architecture provide more available habitat, these variables have been considered as a good proxy of harvestmen diversity in studies at upland Amazonian forests (Colmenares *et al*., 2016, Porto *et al*., 2016, Tourinho *et al*., 2014).

Additionally, habitat quantity and quality have been equally proposed as robust predictors of harvestmen species composition in a fragmented Atlantic forest landscape (Bragagnolo *et al*., 2007), providing more evidence of the importance of habitat quality in maintaining tropical harvestmen assemblages. Fallen woody stems are often considered as one of the most likely microhabitats to contain harvestmen, especially aggregations during diurnal periods (Pinto-da-Rocha *et al*. 2007), yet the associations between harvestmen assemblages and this particular microhabitat have not been properly quantified in tropical forests. In fact, higher density of fallen woody stems may increase habitats favoured by harvestmen, providing ideal microclimatic conditions of humidity and temperature required for the development of some species (Mestre & Pinto-da-Rocha, 2004; Curtis & Machado, 2007; Proud *et al*., 2012). Notably, this was an important variable structuring both harvestmen species composition and abundance across the Balbina islands. A positive association between harvestmen assemblages and fallen logs have not always been detected in undisturbed tropical forests, but this microhabitat type is often associated with some specific harvestmen species in Amazonian forests (Colmenares *et al*., 2016).

The amount of landscape-scale forest cover was another variable retained in the best model influencing havestmen composition on Balbina islands. This variable was also included in the best model affecting patterns of abundance. The role of total amount of surrounding suitable habitat area is widely recognized in determining local patterns of diversity (Andrén, 1994), and has been considered as a good proxy of habitat loss (Fahrig, 2003, 2013). Indeed, several studies in the Atlantic forest has showed the paramount importance of forest cover in predicting abundance patterns of insects (Morante-Filho *et al*., 2016) and vertebrates (Lira *et al*., 2012; Morante-Filho *et al*., 2015)

The amount of surrounding forest cover, number of palm and number of trees, fallen woody stems and island size explained harvestmen abundance patterns at the Balbina islands. Thus, larger islands harbouring higher forest basal area, particularly arborescent palms, are necessary to sustain larger populations of harvestmen species on islands. In fact, tree microhabitats harboured the most diverse and abundant harvestmen assemblages, and was the only variable positively associated with harvestmen abundances in a Central Amazonian forest site distant ∼190 km from Balbina (Colmenares *et al*., 2016).

Overall harvestmen abundance declined in response to local, patch and landscape-scale variables on surveyed islands. For instance, number of trees was a strong predictor of harvestmen abundance patterns. Arboreal trees are highly sensitive to forest fragmentation, with many trees species becoming rapidly extirpated from small forest remnants (Silva & Tabarelli 2000; Laurance *et al*. 2006; Tabarelli, Peres & Melo 2012). This effect is potentially stronger on islands created by hydroelectric dams, with a rapid decay in the floristic diversity detected on small Balbina islands (Benchimol & Peres, 2015b). Fallen woody stems on small Balbina islands – where tree habitat diversity has been severely reduced – may therefore function as important alternative microhabitats for harvestmen assemblages. Therefore, most islands created by the Balbina dam are likely unfavourable for the long-term persistence of viable harvestmen populations, which may lead to local extinctions, creating a set of species-poor islands. On the other hand, species associated with fallen woody stems may benefit from tree mortality induced by high rates of windfalls on small islands (Benchimol & Peres 2015b), thereby becoming highly successful species in this archipelago (two Manaosbiidae and two *Stygnus*-Stygnidae species).

The depauperate tree species composition on most Balbina islands has been detected in a previous study showing that the rapid decay in tree diversity across the archipelago is promoted by edge effects, including edge-related fires and wind-blown tree falls (Benchimol & Peres, 2015b). Although the influence of habitat quantity and structure over the abundance and spatial distribution of harvestmen has not been sufficiently explored, harvestmen have a close relationship with vegetation structure, mostly trees and palms, and microclimate which create environmental gradients affecting their species composition (Colmenares *et al*., 2016). Islands containing higher densities of mature trees and palms harbour distinct assemblages and provide the appropriate microclimatic conditions to retain harvestmen species. Harvestmen and other arachnids, such as spiders, are associated with tree bole surface area, particularly those providing the appropriate bark properties, which are important niches for resting, foraging (Southwood, 1978; Pinzón & Spence, 2010), and appropriate substrates for daytime camouflage and web attachment (Messas *et al*., 2014).

### Species-specific responses to landscape insularization

According to several other studies, the diversity of harvestmen recorded on the islands was low (Pinto-da-Rocha & Bonaldo, 2006; Bonaldo *et al*., 2009; Colmenares *et al*., 2016; Tourinho *et al*., 2014, 2018, Porto *et al*., 2016). The small number of species detected per island also reinforces diversity impoverishment in forest fragments, as suggested for the Atlantic Forest (Bragagnolo *et al*., 2007) and temperate forests (Černecká. et al., 2017, Mihál & Černecká, 2017). Indeed, more than half of the sampled islands exhibited only one to five species, whereas mainland sites retained 11-20 species, exhibiting a similar pattern obtained in other Amazonian sites (Tourinho *et al*. 2011, 2014, 2018, Porto *et al*. 2016, Bonaldo et. al, 2009). The intense sampling effort applied in this study - 20 islands (4 replicates each) - is also an indicative of islands impoverishment (160 hours sampling). As an example, species richness recovered in only 6 controls in mainland areas, with a much inferior effort (48 h) rendered 22 species which may be considered comparable to other areas in the Amazon, and new additions in numbers of sites sampled would certainly increase the number of species sampled as well.

Interestingly, the number of exclusive species retained on islands (9) was not as low as expected, but still lower than the number of species exclusively recorded at mainland forest sites, even if they had been allocated a lower sampling effort (13). However, looking into these results we note that exclusive species sampled on islands are minute harvestmen species and/or specimens, except one. They are members of Agoristenidae, tiny Stygnidae genera, but mostly members of families of Zalmoxoidea and Samooidea (Fig. 2) that inhabit leaf litter layers and other cryptic environments in the Amazon forest, for which little collecting effort had been invested in this study. Given that we sampled few individuals occasionally, mostly singletons or doubletons, the lack of records in mainland sites can also be an artefact, so we would expect to enhance the records of these specimens sifting leaf litter in mainland sites, as happened before in other Amazonian sites (Bonaldo *et al*. 2009, Tourinho *et al*. 2014).

Conversely, Cosmetidae and Sclerosomatidae species seem to be highly responsive to forest alteration and fragmentation. We did not record any of the six mainland Sclerosomatidae species on islands, and from the four species of Cosmetidae, we sampled just 7 specimens of *Eucynortella aff. duapunctata* Goodnight & Goodnight, 1943 on only five islands (Fig. 2), contrasting with 47 specimens sampled across all 5 mainland sites. Those results are quite uncommon because *E. aff. duapunctata* followed by other species of the same genus and at least one or two species of Gagrellinae (Sclerosomatidae) are the most common, widespread and abundant species (Tourinho *et al*. 2014, 2018, Porto *et al*., 2016). Additionally, Cosmetidae, Sclerosomatidae are the top most diverse and abundant species/families in virtually all investigated sites in the Amazon (Bonaldo *et al*., 2009, Tourinho *et al*., 2018, Tourinho *et al*., 2014, Porto *et. al.*, 2016) and our results in mainland followed this pattern as expected in undisturbed areas.

The majority of harvestmen species are highly endemic and have poor disperser skills, tending to move a few dozen meters during their lives (Mestre & Pinto-da-Rocha, 2004, Pinto-da-Rocha *et al*., 2007). Exceptions are some Eupnoi species, such as species of Sclerosomatidae, which exhibited higher vagility. Species of harvestmen are generally predators, and most of the species of Sclerosomatidae observed in the Amazon were predating on small insects (e.g. flies and beetles, Tourinho person. comm.), being more susceptible to habitat and other environmental structure and size than other trophic groups (Srivastava *et al*. 2008, Villanueva-Bonilla *et al*. 2017). Possibly they possess a closer relationship with the structure and architecture of some specific species of trees, palms and/or other plants in the Amazon forests, and because of their physiology and strong associations with plant microhabitat, it might be probably difficult for harvestmen species from forested environments in the Amazon to succeed in colonizing highly disturbed island areas in Balbina lake.

### Conclusions

Arthropods play vital roles in various ecosystem functions, and are widely considered as good indicators of overall biodiversity and forest integrity (Maleque *et al*., 2006). Among the Arachnida, harvestmen comprise the third most diverse order (Shultz & Pinto-da-Rocha, 2007), exhibiting high sensitivity to habitat reduction and edge effects (Bragagnolo *et al*. 2007). Our study shows that forest loss significantly affected the species composition of harvestmen assemblages across islands created by a major hydroelectric dam, following 20 years of isolation. Although total species richness did not respond to local, patch and landscape variables, the assemblage structure of harvestmen was severely affected by landscape-scale forest amount which was probably affected by the rapid decay in tree diversity on most small forest islands in this reservoir (Benchimol & Peres, 2015b). This result demonstrates that harvestmen species composition exhibits a turnover pattern among islands mostly driven by landscape forest amount, therefore indicating that both islands embedded within greater and lower amount of forest cover are important to ensure high diversity of faunal groups. Although we were unable to classify species as generalist and specialist groups, it is expected that islands surrounded by large amount of forest cover are harboring more specialist groups (see Morante-Filho *et al*., 2015). Thus, islands surrounded by high amount of forest must be maintained to preserve sensitive species. We also presume that the rapid decay in tree diversity on most small forest islands in this reservoir (Benchimol & Peres, 2015b), which in turn directly affects habitat quality, is affecting harvestmen species composition. Given the pivotal importance of microhabitat quality for harvestmen (Bragagnolo *et al*., 2007), we expect that a reduction in environmental heterogeneity is directly impacting several harvestmen species. Observed shifts in community structure can potentially lead to an impoverished forest functioning and integrity, especially because changes in tree diversity in insular forests at Balbina are expected to be aggravated in the future (Benchimol & Peres 2015b). We therefore provide additional evidence of the detrimental ecological consequences of dam impoundments on biodiversity, emphasizing that dams are far from the often hailed green sources of energy (Gibson *et al*., 2017). For existing dams, we recommend that well-preserved and large forest islands, surrounded by large amounts of forest cover, should be protected in the long-term. This can buffer the effects of natural forest disturbance and preclude rapid decay in tree communities, contributing therefore to maintain high diversity of several faunal groups, including harvestmen and other arthropods.

## Acknowledgments

Adriano Kury provided key assistance with harvestmen type specimens, Pio Colmenares identified Laniatores specimens and tabulated species data; their support have made this work possible. Regiane Saturnino kindly made available local environmental data collected for her master thesis during arachnids sampling across the Balbina islands. We thank Ricardo Vicente and Thiago Izzo for providing some of the R scripts used in this paper to run some of the analysis. This study was supported by Beca Program, from International Institute of Education of Brazil Programa Beca #B/2005/02/BMP/14, Reserva Biológica do Uatumã and a PNPD grant from the Coordination of Improvement of Higher Education Personnel - CAPES to ALT.

## Authors Contribution

ALT collected data, performed the spatial correlation analysis and wrote the first drafts of the manuscript; ALT and WP identified species, DST ran most of the analysis, made figures, tables and described these procedures in Methods section. MB ran analysis and actively participated in the subsequent writing rounds. ALT, DST and MB conceptualized the study. CP detailed checked the results. All the authors discussed the results, read and equally contributed to the subsequent versions of the manuscript.

---

**Table S1.** Number of individuals and species of harvestmen (Arachnida: Opiliones) sampled across 20 islands and 5 continuous forest sites across the Balbina Hydroelectric reservoir, Central Brazilian Amazonia.

| **Site name** | **Area**<br>**(ha)** | **UTM (X)** | **UTM (Y)** | **Number of**<br>**species** | **Number of**<br>**individuals** |
| --- | --- | --- | --- | --- | --- |
| Angelin | 67.44 | 240659 | 9801730 | 4 | 9 |
| Cachoeira | 11.84 | 236641 | 9793981 | 5 | 20 |
| Cipoal | 213.66 | 190871 | 9810812 | 5 | 17 |
| Copa | 19.20 | 229635 | 9798056 | 4 | 13 |
| Folharal | 102.30 | 192046 | 9806910 | 6 | 17 |
| Galo de briga | 190.84 | 197368 | 9807612 | 3 | 9 |
| Gemeas | 111.09 | 203037 | 9814880 | 6 | 38 |
| Grande | 95.61 | 204026 | 9806251 | 6 | 48 |
| Igarapé | 76.20 | 236013 | 9805808 | 4 | 10 |
| Josue | 146.00 | 205982 | 9801462 | 6 | 24 |
| Manoel gato | 194.59 | 223583 | 9802328 | 6 | 18 |
| Massaranduba | 17.33 | 201171 | 9802527 | 7 | 25 |
| Mucura | 22.57 | 229798 | 9800932 | 1 | 1 |
| Neto | 41.59 | 237861 | 9796638 | 4 | 5 |
| Palhal 1 | 20.78 | 237245 | 9800112 | 2 | 5 |
| Palhal 2 | 25.81 | 227557 | 9801956 | 6 | 16 |
| Pontal | 126.01 | 201747 | 9797850 | 8 | 27 |
| Santa Luzia | 218.53 | 228363 | 9808143 | 3 | 4 |
| Serrinha | 50.09 | 231033 | 9792680 | 2 | 2 |
| U | 79.83 | 204181 | 9799600 | 10 | 49 |
| Mainland 1 | ∞ | 251524 | 9801136 | 14 | 89 |
| Mainland 2 | ∞ | 250493 | 9801920 | 11 | 55 |
| Mainland 3 | ∞ | 247494 | 9800939 | 15 | 67 |
| Mainland 4 | ∞ | 251567 | 9798861 | 20 | 168 |
| Mainland 5 | ∞ | 251564 | 9797900 | 11 | 47 |
| TOTAL |  |  |  | 31 | 783 |

